# A global synthesis of phenological responses to climate change

**DOI:** 10.1101/164806

**Authors:** Jeremy M. Cohen, Marc J. Lajeunesse, Jason R. Rohr

## Abstract

Phenology, or the timing of seasonal activities, is shifting with climate change, resulting in disruptions to the timing of migration and breeding and in emerging asynchronies between interacting species^1-5^. Recent syntheses have concluded that trophic level^1^, latitude^6^, and how phenological responses are measured^7^ are key to determining the strength of phenological responses to climate change. However, despite these insights, researchers still lack a comprehensive framework that can predict responses to climate change globally and across diverse taxa. For example, little is known about whether phenological shifts are driven by different climatic factors across regions or which ecologically important species characteristics (e.g., body size) predict the strength of phenological responses. Here, we address these questions by synthesizing hundreds of published time series of animal phenology from across the planet. We find that temperature drives phenological responses at mid-latitudes, but precipitation is more important at lower latitudes, likely because these climate factors often drive seasonality in each of these regions. Body size is also negatively associated with the strength of phenological shift, suggesting emerging asynchronies between interacting species that differ in size, such as hosts and ectoparasites and predators and prey. Finally, although there are many compelling biological explanations for spring phenological delays, some examples of delays are associated with short annual records prone to sampling error. As climate change intensifies, our findings arm biologists with predictions concerning which climatic variables and organismal traits drive phenological shifts.

---

Global climate change has significant ecological consequences^4,8^ and perhaps the best-studied are advancements in the timing of seasonal activities, or phenology, of organisms^1-3,5,7,9-13^. Understanding the factors influencing phenological shifts are critical because these shifts can impact the fitness of organisms by altering the availability of resources^2-4^. In addition, phenological shifts can cause species declines by generating asynchronies or “mismatches” between plants and pollinators^12^, plants and herbivores^14^, migrant birds and their prey^11^ or floral resources^15^, and hosts and parasites^16^. Several recent syntheses have made significant inroads to understanding how the phenology of species is shifting with climate change^1,5-7,13^. For example, primary consumers were demonstrated to be shifting their phenology faster than other species in the U.K.^1^, species are shifting their phenology faster in spring than in autumn in China^5^, and the strength of phenological responses to climate change is dependent on the way responses are measured e.g., type of behavior observed or number of observations ^7^.

Despite these insights, several critical knowledge gaps preclude accurate predictions on the sensitivity of organisms to climate change on a global level. First, although many phonological syntheses assume climate change as an important driver, few explicitly test for effects of climate (but see^1,5,6^), and among those that do, climate data that has been standardized among studies is rarely used to confirm the link between changes in phenology and climate. As a consequence, it remains unclear which climatic variables, such as temperature or precipitation, are driving shifts in phenology, and whether the broad geographical heterogeneity in these climate variables impacts the power of these variables to explain and predict ecological trends. Second, recent syntheses have relied on country-level data, and no synthesis in over a decade has addressed phenological responses to climate change across the globe. Global analyses greatly improve sample sizes and statistical power to detect drivers of phenological shifts. In addition, local-scale syntheses may miss important broad-scale spatial heterogeneity in climate factors. For example, global syntheses are critical to test broad-scale latitudinal hypotheses about phenological shifts, such as the hypothesis that climatic factors driving seasonality across latitudes also drive phenological changes. Third, it is unclear why some species show delayed spring phenologies despite an overall trend towards advancement^10,17^. Finally, it is also unclear whether certain ecologically important characteristics of organisms are predictive of strong phonological responses. For example, body size may be an important factor determining the magnitude of phenological response to climate change because smaller organisms acclimate more quickly to changing conditions than larger organisms^18^. In addition, ectotherms may exhibit stronger phenological responses than endotherms because they cannot thermoregulate independent of their environments and are therefore more sensitive to changes in environmental conditions. Because of these knowledge gaps, a general global framework is still missing for predicting the direction and magnitude of phenological shifts based on ecological context and organismal traits.

To address these gaps, we conducted a global synthesis of phenological time series of animals from 127 studies (Table S1; Table S2), spanning five continents and 15 classes of animals including insects, mammals, reptiles, and birds. We focused on spring phonological events in animals because phenological responses to climate change in plants have recently been synthesized^19^, some of our primary questions could only be answered using animal data, and the evidence for advancement in animal phenology is more conflicting and controversial than it is for plants^9^ (see Supplement). Here, we synthesize the multivariate effects of climate change on phenology, as well as test predictors of this complex phenomenon (e.g., latitude, endo – or ectothermy), with a unique meta-analysis approach that jointly models phenological shifts, the effects of climate on phenology, and climate change (the 50 year correlation between climate and year) using a trivariate mixed-effects model^20,21^ (see Extended Data Fig. 1; see Methods). Unlike previous univariate meta-analyses that strictly synthesize phenological shifts^2,3^, our trivariate approach assesses whether phenology is dependent on climate and climate change and whether the magnitude and direction of these relationships is dependent on 10 climate variables (e.g., mean, minimum and maximum temperature, precipitation, snowfall^22^, see Methods). All climate variables were standardized across all time series by accessing a single source of historical point-based climate data (NOAA’s NCDC-3 data^23^) with data that were specific to the region and time of each study, reliably allowing us to identify which aspects of climate were driving phenological shifts. Importantly, this approach facilitated evaluation of whether climate change (the 50 year slope between climate and year), rather than just long-term climate means, was associated with changes in phenology. Further, our trivariate mixed-effects meta-analysis also accounted for dependencies of effects among related taxa due to their shared phylogenetic history^24^ (see Supplementary Code). We were able to compare relationships between phenology and year for 1,011 time series and relationships between phenology, year and climate for a subset of these including 321 time series.

**Figure 1:**
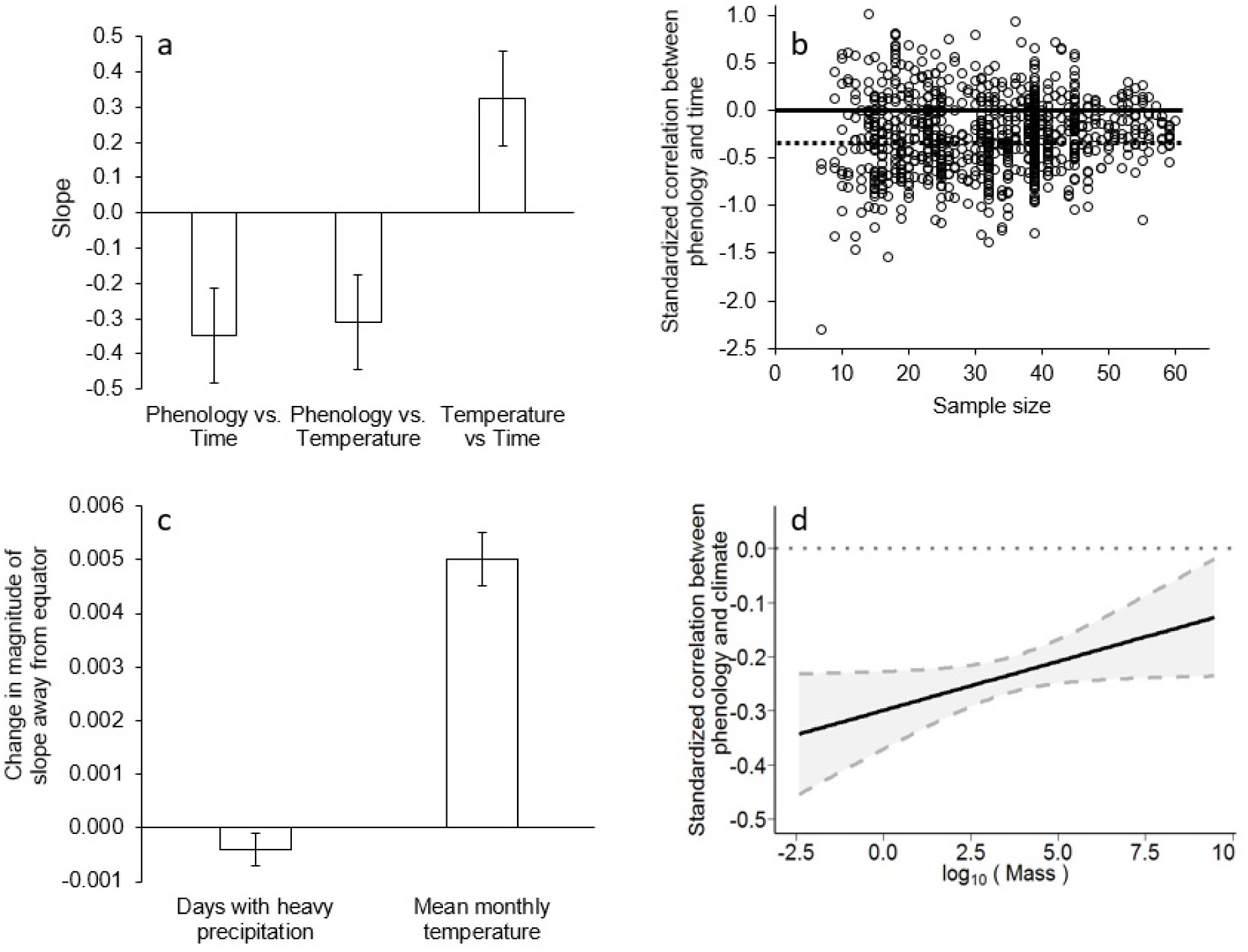
Improving how we understand advancements in phenology due to climate change. (a) Across 1,011 time series, phenology became earlier through time as temperature increased through time and the detrended increases in temperature were negatively correlated with phenology (see Extended Data Fig. 4 for precipitation). Error bars represent SEM. (b) A funnel plot comparing sample sizes (total years in time series) with standardized effect sizes (correlation between phenology and time quantified via Fisher’s Z effect sizes) reveals that studies with small samples sizes have large variation with both the positive and negative shifts, suggesting that species appearing to delay their phenology in spring might sometimes be spurious products of sampling error. The solid line is the zero line and the dotted line represents the grand mean correlation (- 0.349). (c) The slope between temperature and phenology became steeper as the absolute value of latitude (or distance from the equator) increased (bar, p<0.0001), while the slope between rainfall and phenology became less steep (p<0.01). Thus, temperature became more predictive of phenology further from the equator, whereas rainfall became more predictive of phenology closer to the equator, suggesting that the phenology of species is driven by the climatic factor that drives seasonality locally. Error bars represent SEM. (d) The slope between log-transformed body mass and the correlation between phenological date and mean temperature is positive in the trivariate meta-analysis model, indicating that smaller organisms track their phenology with temperature more closely than larger organisms. Data points are not shown to reduce clutter and 95% confidence intervals are provided in gray.

The meta-analysis revealed that, on average, animals have advanced their phenology significantly since 1950 (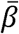, df = 937, *p* = 0.01; Fig. 1a; Table S3), advancing by 2.88 days/decade and 3.08 days/°C. Across all species and sites, mean temperature increased significantly over time (Fig. 1a; Table S4). The meta-analysis also revealed that temperature is closely related to phenological date independent of year, and that phenology is more closely linked with mean temperature in areas with more climate change (Extended Data Fig. 2), suggesting that climate change is indeed the driver of these shifts (Fig. 1a; Table S4). Phenological shifts were not heavily biased by the phylogenetic history of taxa, which accounted for only about 4.5% of the variance (phylogenetic τ**^2^**) between phenology and year, and 0-6% between phenology and climate (Tables S3-8). Between-study variance accounted for 8-9% of the total variance accounted for in all models (Tables S3-8).

**Figure 2:**
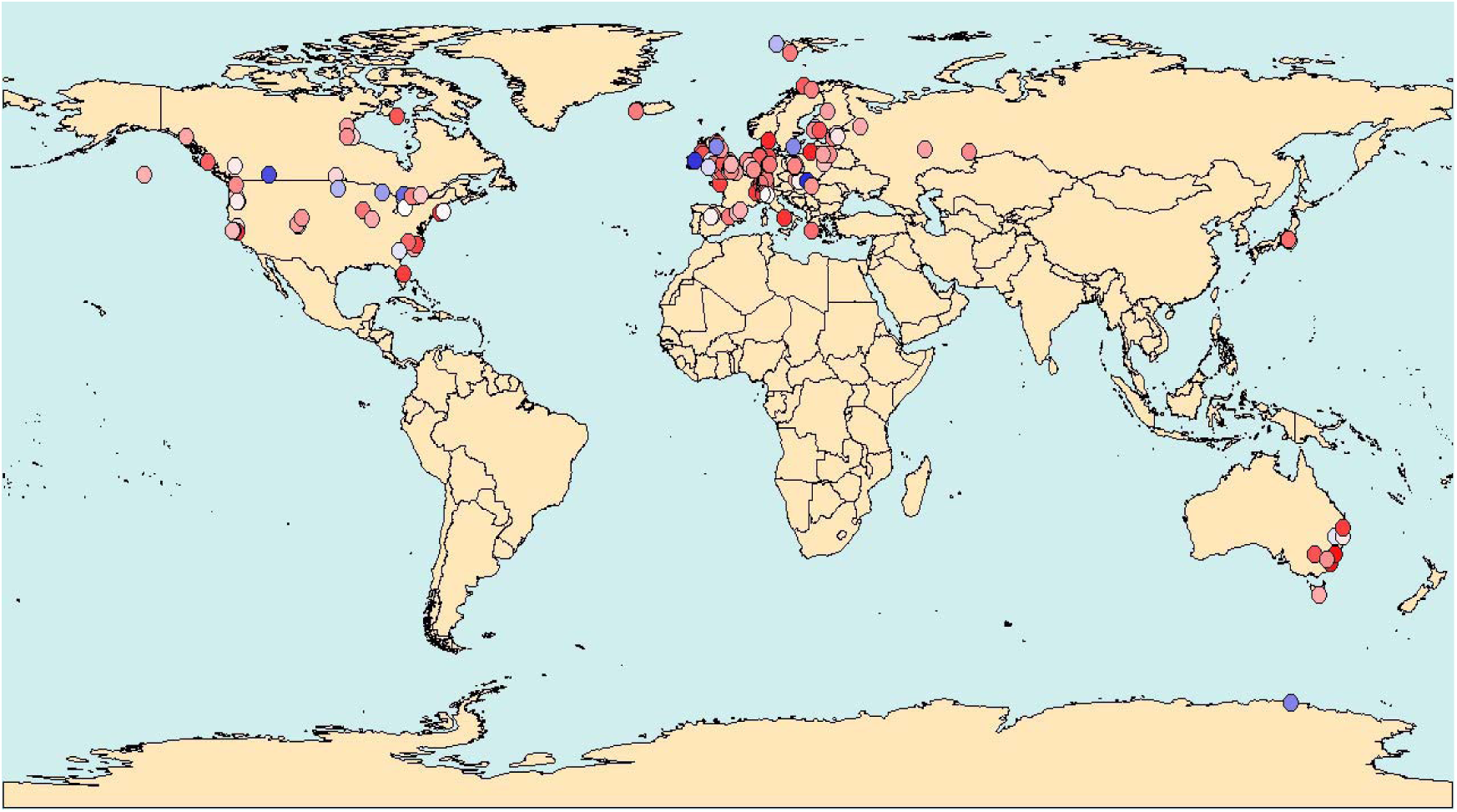
The uneven global distribution of published studies exploring the phenology of animals. There are hundreds of published phenology time series from North America and Europe, but much less is known about phenology on the other five continents with particularly large gaps in the tropics and marine systems. Red points indicate advancements in phenology over time and blue points indicate delays. The strength of the color indicates the magnitude of the relationship between phenology and time (as quantified with a Fisher’s Z effect size).

The direction of phenological shifts may differ among taxa, with some species showing delays rather than advances of spring phenology^5,10,13,17,19^ — such as delays in seabird egg laying as a consequence of reduced sea ice^10^ or delays in phenology (e.g., flowering) after short winters that fail to induce vernalization^17^. To test whether a similar phenomenon might be responsible for phenological delays (positive relationships between phenological date and year), we examined whether the magnitude of the delay could be predicted by the increase in winter temperatures (defined here as the relationship between year and average temperature during the year’s three coolest consecutive months), controlling for latitude. We found no support for the hypothesis that winter temperatures predicted phenological delays, instead finding that they predicted advancements (β =-0.296, df = 321, *p*<0.001 in models with all time series) or were not predictive (β =-0.125, df = 68, *p* = 0.32 among time-series with delays only). In fact, winter temperatures were positively correlated with spring temperatures that are well documented to drive phenological advancements (β = 0.298, df = 321, *p*<0.0001 for all time series, β = 0.202, df = 68, *p* = 0.03 among delays). Alternatively, many apparent spring delays might be sampling artifacts of short annual records. Indeed, a funnel plot revealed that many studies based on short time series (e.g., small sample sizes) had both delays and strong advances in phenology, but when sample sizes were large, phenology advanced more uniformly (Flinger-Killeen test for homoscedasticity: ?^*2*^ = 112.72, *p*<0.0001; Fig. 1b; see Extended Data Fig. 3 for comparisons of effect sizes with variance). In addition, there was no evidence of funnel plot asymmetry (Egger’s test: *z* =-0.724, *p* = 0.47), suggesting that phenological delays are appropriately represented in our dataset. While this result does not exclude true and biologically relevant spring delays in phenology (see examples above), it suggests that reports of delays are likely sensitive to sampling error; in fact, duration of time series has previously been found to influence observed phenological trends in marine species^7^. These findings also indicate that previous phenological syntheses^1-3^ may have underestimated phenological advancements since they did not statistically account for differences among studies emerging via sampling error^25^.

**Figure 3:**
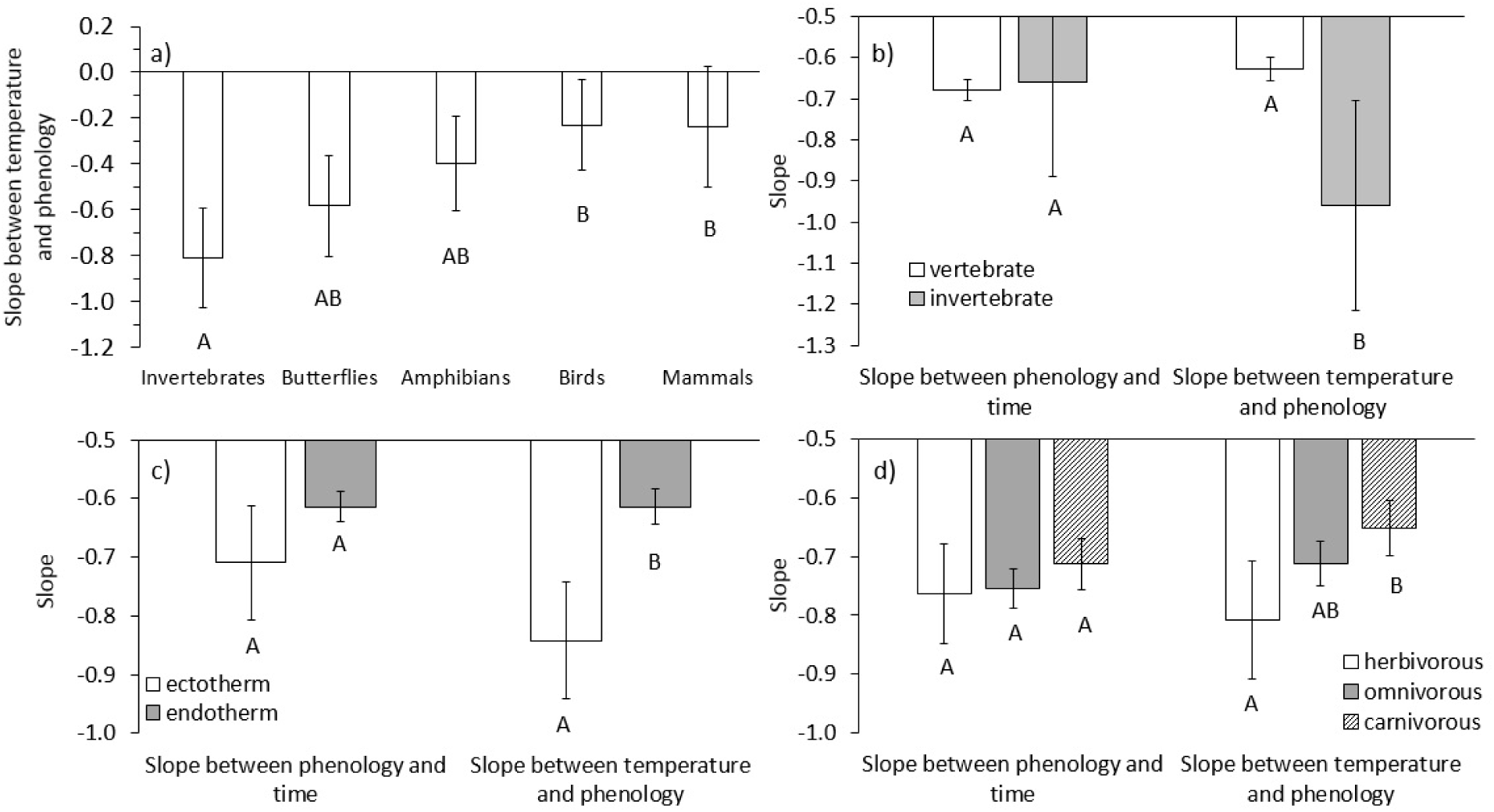
The ability of phenology to track temperature varies among taxonomic classes of animals, ecto-or endothermy, and trophic level. In models including body size as a covariate, (a) smaller taxa, such as (b) invertebrates, and (c) ectotherms tracked temperature closer than larger animals and endotherms. (d) Herbivores had a greater association between temperature and phenology than carnivores, possibly because herbivores were reacting to shifts in plant phenology associated with temperature. We report relationships between phenology and both temperature and time to highlight that even if groups are apparently advancing their phenology at similar rates, they could be responding to changing climates at dissimilar rates if they come from regions experiencing different rates of climate change. Error bars represent SEM for the slope parameters from trivariate mixed-model meta-regressions. Different letters denote statistically significant differences in effect sizes.

We also hypothesized that phenological shifts would be associated with the climatic variables that drive seasonality locally — such as temperature at mid-latitudes (i.e. temperate zones) and precipitation at low latitudes (i.e. tropical and subtropical zones). Moreover, because climate change is resulting in greater changes in temperature than precipitation^26^, we hypothesized greater phenological shifts in temperate than tropical zones. In support of these hypotheses, as absolute value of latitude increased, changes to temperature became more predictive of the magnitude of phenological shifts, and as latitude decreased, precipitation was a stronger predictor of phenology (test for different slopes^27^: *t* = 7.89, df = 1650, *p*<0.0001; Fig. 1c; Table S5). Further, there was a greater increase in temperature than precipitation through time (Extended Data Fig. 4), and the correlation between phenology and temperature in the temperate zones was stronger than the correlation between phenology and precipitation in the tropics (Fig. 1c). These results indicate that different climatic variables are triggering phenology in temperate and tropical regions. While past syntheses have hypothesized that species should shift their phenology faster at higher latitudes in response to greater warming in these regions^2,3,6^, low-latitude species may also be shifting their phenology at high rates in response to changes in rainfall. Given that the majority of phenological studies are from northern temperate climates^7^ (especially North America and Europe; Fig. 2), and emphasize temperature over precipitation, additional phenological time series from low latitudes are needed to quantify the full effects of precipitation shifts on tropical phenology. However, the effects of precipitation on phenology may be less closely associated with latitude than the effects of temperature simply because latitude is more strongly correlated with temperature than precipitation.

Given that temperature and precipitation drive phenology unequally across the globe and particular taxa exhibit differential sensitivities to extreme temperatures and moisture levels, we hypothesized that the phenology of specific taxonomic groups might be more strongly associated with temperature than precipitation. For example, we expected amphibians to respond to precipitation more strongly than any other taxonomic group because of their considerable reliance on moist conditions for survival and reproduction. Across all taxa synthesized, phenology was associated more strongly with temperature than with precipitation (temperature,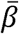=-0.310, df=1579, *p*=0.02;precipitation, 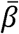=-0.054, df=1579, *p*=0.54; Extended Data Fig. 5; Table S4), and different components of temperature (mean, minimum and maximum) did not significantly differ from one another at predicting phenology. As predicted, amphibians exhibited the strongest association between precipitation and phenology among all taxa (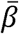; Extended Data Fig. 5b; Table S6).

Next, we sought to identify general ecologically-important characteristics of taxa that might predict the strength of phenological responses to climate change. Here, we hypothesized that ectotherms and smaller organisms should be more sensitive to shifts in climate than endotherms and larger organisms (because thermal inertia is positively associated with body size^18,28^). Indeed, body size was a significant negative predictor of phenological shifts alone (β =-0.0221, df = 921, *p*<0.01; Table S7) and as a covariate alongside other species traits in most statistical models (Fig. 1d; Table S7). Among taxa, invertebrate groups tracked their phenology to temperature more closely than birds and mammals (Fig. 3a; Table S7), and as a whole, invertebrates tended to track temperature better than vertebrates (Fig. 3b; Table S7). As predicted, the phenology of ectotherms was more strongly correlated with temperature than the phenology of endotherms (Fig. 3c; Table S7). Finally, herbivore phenologies tracked temperature more closely than carnivore phenology (Fig. 3d; Table S7) when accounting for body size, possibly because herbivores are also responding to shifts in the timing of plant phenology^29^, supporting conclusions by Thackeray *et al.* based on phenological shifts in the U.K. ^1^. Additionally, we did not observe a difference between the phenological responses of terrestrial and aquatic species (Extended Data Fig. 6; Table S7), although there are admittedly few aquatic species in the dataset (18 total) and all are marine.

Finally, we posited that the type of phenological responses, such as peak seasonal abundance, arrival (migration), and breeding/rearing (calling, nesting, laying, hatching, or weaning), may differ in their sensitivities to climate change, as recently concluded by synthesis on marine systems^7^. We predicted that arrival would be least correlated with climatic factors because migrants are likely reacting to climatic conditions where they left from rather than conditions where they are arriving^30^. Phenological responses related to arrival tracked climate the most poorly (Extended Data Fig. 7; Table S8), and those based on peak abundance tracked temperature changes the most closely — possibly because peak abundance is more often documented with smaller invertebrates that phenologically respond strongly to climate. Unfortunately because there are very few phenological time series from equatorial regions, and arriving species often come from multiple departure locations, we could not test whether the timing of departures for spring migrations tracked temperature better than arrivals (but see^31^).

Our findings add to the growing evidence of direct ecological consequences of climate change on ecological systems and provide strong evidence linking climate change to phenological shifts. Our synthesis unveiled previously unidentified generality in the phenological responses of organisms to climate, indicating that the phenology of species at high and low latitudes most strongly respond to temperature and precipitation, respectively, and thus different components of climate drive phenology in different regions of the globe. We also found that different taxa respond to the same climatic signals but do so at different rates, and that the strength of these phenological shifts is predictable based on two easily measured traits, thermoregulation and body size. As climate change intensifies in the next century, our results suggest that advances in phenology are likely to become more exaggerated, potentially further desynchronizing interactions between species that vary considerably in their body sizes, such as mutualistic, predator – prey, and host – parasite interactions. However, the synthesis presented here now arms climate biologists with knowledge regarding the specific components of climate and the traits of interacting species that can drive phenological shifts, providing new opportunities to forecast mismatches and mitigate their adverse effects.

## Materials and methods

### Literature survey and data requirements

We conducted a literature search in September 2012 on Web of Science for the term “phenology AND climate” within the following fields: environmental sciences and ecology, zoology, developmental biology, reproductive biology, life sciences (other), entomology, behavioral sciences, physiology, biodiversity and conservation, fisheries, evolutionary biology, parasitology, marine and freshwater biology, infectious diseases, and oceanography. This search generated 6,989 studies which were examined for phenological time series. References in these papers and the USA National Phenology Network (usanpn.org) database were also examined for time series. Time series were not used if they (1) contained data from a span of <10 years; (2) contained data for fewer than seven individual years; (3) described autumn migrations; or (4) described data that were redundant with data we had already compiled from another paper. We also eliminated raw data from before 1950, because this is considered to be before significant global climate change^32^. Our exclusion criteria are similar to those from previous meta-analyses^2,3^.

### Data extractions

We extracted raw time series data from figures plotting day of year of phenological event (including date of first or median arrival, first calling, nesting, laying, peak abundance, oestrus, or weaning) against year using Datathief III Version 1.6 (© Bas Tummers). Correlation coefficients between phenological date and year, standard errors or surrogates, and slopes were also calculated for each time series when they were not reported in the original text (All analyses were conducted in R 3.1.0; stats package, glm function). Correlation coefficients (*r*) and standard deviations were available for 1,011 of these time series (representing 127 studies) which were used in the meta-analysis examining the relationship between phenology and year. Approximately 400 time series from about 100 papers provided raw data and were used in the meta-analyses examining the relationships between phenology, year, and climate (the actual numbers varied between different climate variables because some variables were not available at certain geographic locations). Sampling variances (used as weights) were derived from all correlation coefficients, and coefficients and variances were standardized using Fisher’s z-transformation before all meta-analysis modeling.

### External climate data

Climate data were from the NOAA National Climatic Data Center (NCDC; <u>www.ncdc.noaa.gov</u>) worldwide database of “monthly observational data” corresponding to the nearest location (within 100km) and full time span of every time series that provided raw data and geographic coordinates. Ten climate variables were used in our meta-analysis (see Extended Data Fig. 6), and they generally were related to temperature or precipitation. Climate variables were used individually in models instead of as covariates (see below). Yearly averages of climate variables were compiled for all variables in all locations and for the years in all time series only when data were available for all 12 months. Within each time series, correlation coefficients and standard errors were compiled for all correlations between all annual climate variables year, all climate variables and phenology, and phenology and year (stats package, glm function). We did not have any climate data for marine species and did not include these time series in any analyses testing the effects of climate.

### Independent fixed-effects variables

Independent variables collected for each time series included taxonomic classification of the focal species, absolute value of latitude, elevation, form of thermoregulation (ectothermy or endothermy), trophic level, habitat (terrestrial or marine), country (to control for geography), log-transformed body mass (see Supplementary Methods) and type of phenological event (endpoint measured). Taxonomic classification was assessed to the class level. Elevation specific to the locations where time series were observed was extracted from Worldclim elevation rasters (<u>www.worldclim.org</u>) (raster package, extract function). Trophic levels were assigned categorically as “herbivore”, “omnivore”, or “carnivore”. If a species typically eats plants and animals it was designated an omnivore, but if it mostly relies on either prey or plants and only occasionally ate the other, it was assigned to “carnivore” or “herbivore” respectively. Phenological events were categorized as either “arrival” (migrations), “breeding/rearing” (calling, nesting, laying, hatching, or weaning), or “peak abundance” (peak population abundance).

### Meta-analysis models

A trivariate mixed-effects meta-analysis was used to analyze three effect sizes per study that jointly quantify the pairwise relationships among phenology, time, and a single climate variable. Preserving the trivariate structure of effect sizes has the advantage of accounting for the correlations within the three non-independent effect sizes (because of sampling variability and covariances), while also explicitly accounting for any existing correlations among these three effect size groups (via a multivariate random-effects model). Our overall model had a hierarchical structure in which we modeled the sampling variances and covariances among the three effect sizes (within-study weighting to account for study sampling error; see Supplementary Methods), between-study random-effects for each effect size triplicate that were allowed to be correlated but differ among groups (i.e., a multivariate version of the between-study variance component typically included in traditional random-effects meta-analysis), and finally an unstructured random-effect modeling the phylogenetic correlations among taxa (see Supplementary Methods and Code). For all models, the *rma.mv* function from the R package *metafor* was used (see Supplement), with the variance-covariance matrix as the variance-covariance matrix of the sampling errors, and all random effects (trivariate between-study variances, and phylogenetic) were based on restricted maximum likelihood estimator using a nlminb numerical optimizer. However, we did not include phylogenetic random-effects in our analyses of the relationship between phenology and body size because phylogeny and body size are highly correlated and thus controlling for phylogeny also indirectly eliminates much of the body size variation (see Supplement). Please see Supplementary Code for the R script used in these analyses.

## Acknowledgements

We thank N. Argento and C. Gionet for assistance extracting data from studies, T. James for assistance compiling references, C. Parmesan for helpful discussions on vernalization and phenological meta-analyses in general, and D. Civitello, B. Delius, N. Halstead, S. Knutie, K. Nguyen, N. Ortega, B. Roznik, E. Sauer and S. Young for manuscript comments. This research was supported by grants from the National Science Foundation to M.J.L (DBI-1262545, DEB-1451031) and J.R.R. (EF-1241889, DEB-1518681), and National Institutes of Health (R01GM109499, R01TW010286), US Department of Agriculture (NRI 2006-01370, 2009-35102-0543), and US Environmental Protection Agency (CAREER 83518801) to J.R.R.

## Author information

Reprints and permissions information is available at <u>www.nature.com/reprints</u>. The authors declare no competing financial interest. Correspondence and requests for materials should be addressed to J.M.C.

